# Epigenetic markers of middle-age: non-linear DNA methylation changes with aging in humans

**DOI:** 10.1101/2025.08.14.670237

**Authors:** Akiyoshi Shimura, Kyosuke Yamanishi, Takaya Ishii, Tomoteru Seki, Tsuyoshi Nishiguchi, Bun Aoyama, Therese Santiago, Varun B Dwaraka, Ryan Smith, Gen Shinozaki

## Abstract

**Background:** Human DNA is known to exhibit an overall tendency toward demethylation with aging. However, assuming a simple linear relationship between DNA methylation and age does not align with the phenotype of human development and the aging process. This study aimed to investigate the existence of DNA methylation patterns with peaks or troughs at specific ages in addition to simple linear changes.

**Methods:** A large-scale dataset of genome-wide DNA methylation data from 10,420 individuals was analyzed. Hierarchical multiple regression models were applied to detect patterns of the association between age and DNA methylation: linear increase, linear decrease, U-shaped curve, and inverse U-shaped curve.

**Results:** Among the 864,627 CpG sites analyzed, 8.4% exhibited an increase in DNA methylation with age, 23.9% showed a decrease, and 5.5% were better explained by a quadratic model (P < 5.7815×10 ). Within the non-linear subset, inverse U-shaped CpG sites peaking in methylation during middle age were predominant. Genes exhibiting quadratic association patterns between DNA methylation and age, and those linked to diseases with common onset during middle age, were also detected.

**Conclusions:** Non-linear age-related DNA methylation patterns, with peaks or troughs occurring at specific ages, were detected. This suggests that humans do not simply age linearly, but that programmed mechanisms or cascade-like processes may exist to promote or suppress the expression of specific genes at certain ages, contributing onset of certain diseases at specific timings.

## Introduction

DNA methylation is one of the epigenetic processes that regulate gene expression, primarily through the addition of a methyl group to the cytosine bases commonly followed by guanine (CpG: 5’—C—phosphate—G—3’). The addition of methyl groups in the gene promotor region inhibits access of RNA polymerase [1], leading to the repression of gene expression. DNA methylation patterns are known to change with age. In vertebrates, total methylated cytosine levels in DNA tend to decrease with aging [2]. In humans, a previous study investigating DNA methylation across a wide range of blood samples from individuals aged 14 to 94 years with 421 samples reported that approximately 30% of all CpG sites change with age, with 60% of these sites showing a trend toward demethylation as people age [3]. Conversely, methylation tends to increase in promoter regions. These age-related methylation changes are known as "epigenetic drift," characterized by gradual, extensive demethylation of the genome alongside hypermethylation of several promoter-associated CpG islands [4]. However, the mechanisms by which aging induces these changes, leading to aging-associated differentially methylated regions, remain unclear. Potential mechanisms of hypomethylations include age-related changes in DNA methylation and the repair pathways [5], and the impact of oxidative stress [6]. Nevertheless, no comprehensive explanation has been provided for why DNA methylation changes in both positive and negative directions with age.

Despite the unclear mechanisms, robust evidence has shown that specific DNA methylation levels at CpG sites in certain genes change with age. Numerous studies have utilized DNA methylation data to predict chronological age or phenotypic age, with these predictions demonstrating high accuracy [7]. Age estimation based on DNA methylation status is suggested to be more accurate compared to other molecular biological markers [8], and its utility has been proposed in forensic science [9]. On the other hand, a common limitation of previous studies investigating the relationship between age and DNA methylation is that they categorize CpG sites into three groups: those showing increased DNA methylation with age, those where DNA methylation decreases, and those without significant age-related associations. In other words, these studies all assume a linear association between age and DNA methylation. However, biological functions in the human body do not necessarily change linearly with age. For example, obvious non-linear age-related responses in the human body include the secretion of growth hormones [10] and sex hormones [11, 12]. The total secretion of these hormones is low during childhood, peaks during adolescence and early adulthood, and declines in old age, forming an inverse V- or U-shaped relationship with age. Thus, humans experience gene regulation depending on different life stages, from birth, development, growth, reproduction, and aging. It is therefore unreasonable to assume that DNA methylation level and gene expression uniformly increases or decreases linearly throughout the life span. Instead, it is likely that gene activity is enhanced or suppressed in a manner suited to each life stage.

Here, we hypothesize that humans have DNA methylation that exhibits peaks or troughs at different ages in various patterns beyond linear changes. To test this hypothesis, we analyzed age-related DNA methylation changes using a large dataset comprising more than 10,000 individuals from the general population. We classified the changes into five categories: linear increase, linear decrease, inverse U-shaped peaks at certain ages, U-shaped troughs at certain ages, and no significant age association. We then investigated the proportions of these patterns and the associated gene groups (Figure 1). This will allow the identification of genes whose expression is specifically promoted or suppressed at certain ages. Furthermore, it may provide insights into the mechanisms underlying age-related diseases with specific onset periods and thus lead to discovery of novel target for effective intervention. It may also help uncover potential cascades involved in the aging process.

**Figure 1.**
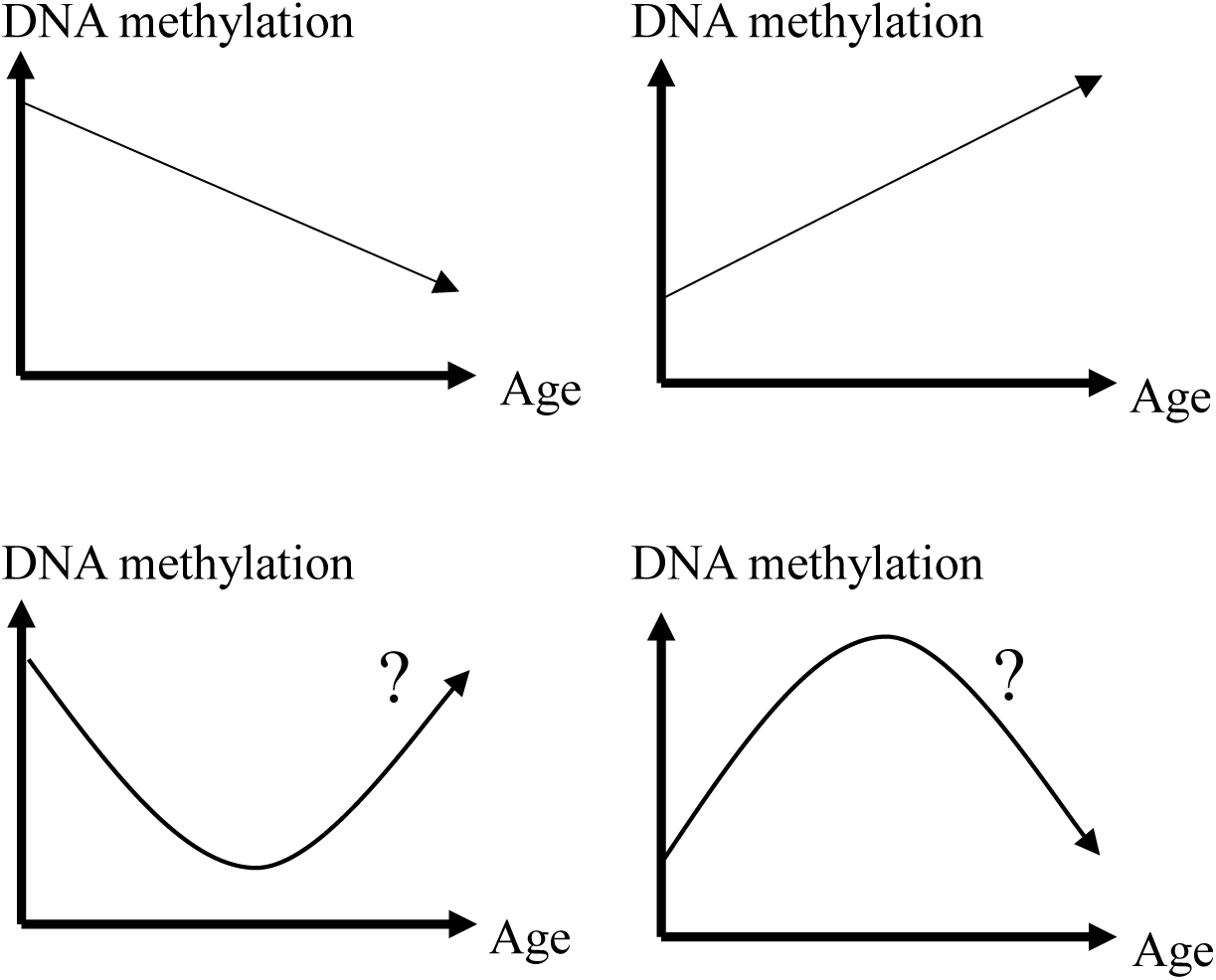
**Explanation of the model**

## Methods

### Dataset

A large-scale, fully anonymized and de-identified genome-wide DNA methylation database was used for the analyses. The dataset, provided by TruDiagnostic Inc. (Lexington, KY, US), includes DNA methylation and demographic data from individuals who consented to the academic use of their data through the epigenetic age measurement service conducted between 2021 and 2022. The data collectors, holders, and analysts are independent of each other. The database contains data from 10,420 individuals, encompassing DNA methylation level data at 864,628 CpG sites obtained using the Illumina Methylation EPIC Bead Chip microarray [13]. The analysis of this dataset was approved by the Institute of Regenerative and Cellular Medicine Institutional Review Board (IRB Approval number: #IRCM-2023-369). This study does not involve interaction with participants or identifiable private information or biospecimens, and it has therefore has been exempted from the Stanford IRB review (Protocol ID: 76650).

### Blood collection and DNA methylation dataset creation

Participants self-collected blood samples using a finger-prick device, which enabled the collection of small amounts of blood from the fingertip. The blood samples were stabilized and shipped to the lab at room temperature. Upon arrival at the laboratory, DNA was extracted and purified from the whole blood samples. After quantification, the DNA underwent bisulfite conversion and was then analyzed using the Illumina EPIC methylation array. To pre-process the array data, we used the *minfi* pipeline [14], and low quality samples and probes were identified using the *qcfilter* function from the *ENmix* package [15], using default parameters. A total of 10,420 individuals passed the quality control (P < 0.05). Finally, β values of all CpG sites were included in the dataset. The composition ratio of white blood cells (WBC) was estimated using EpiDISH [16, 17].

### Statistical analyses

Hierarchical multiple regression analyses were conducted on all 864,628 CpG sites. In the first step, multivariate linear regression analyses were performed with the β values set as the dependent variables, and age (years), sex (male=1, female=0), and estimated white blood cell (WBC) composition (percentage of CD4+ T cells, CD8+ T cells, B cells, NK cells, and monocytes; %neutrophils were used as a reference) were included as independent variables for adjustment. In the second step, the quadratic term of age (age²) was added to the regression model as an independent variable (Figure 2A). Changes in the coefficient of determination (R²) and the significance of these changes were tested. If the age² term and change in R² were significant, the peak or trough age was determined by the quadratic regression model. The peak or trough age can be calculated by dividing the regression coefficient of the age term by -2 times the regression coefficient of the age² term (Figure 2B).

**Figure 2.**
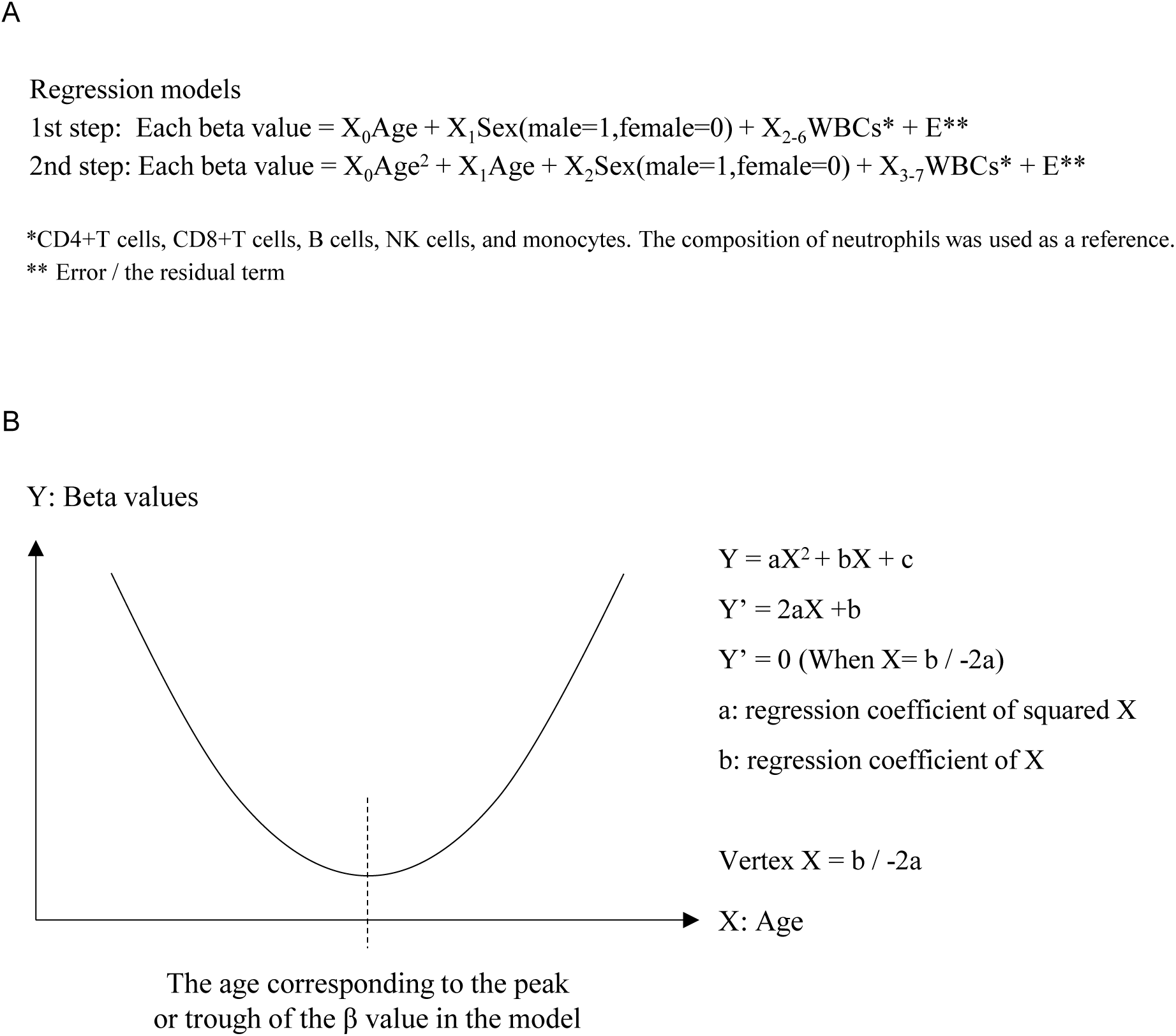
**Explanation of quadratic equation model regression analysis**

Throughout the regression analysis, both regression coefficients, indicating the change in DNA methylation levels (β values) per one-year change in age, and standardized regression coefficients, representing the change in standardized DNA methylation levels per one standard deviation (SD) change in age, were calculated. For β values with large versus small variances, the physiological meanings of changes in β values associated with age may differ even if the absolute change is the same. This difference could be assessed using standardized regression coefficients.

The threshold for statistical significance was set at a 5% P-value of <0.05. For the epigenome-wide significance threshold, considering multiple comparisons, a Bonferroni-corrected threshold of P < 5.7815×10 (=0.05/864,827) was established. This threshold is more conservative than the previously recommended threshold of P < 9×10 for EWAS using the EPIC array [18] and is almost the same as the threshold used in genome-wide association studies (GWAS) involving over a million single nucleotide polymorphisms (SNPs) [19].

### Visualizing

First, several plots were generated for visualization. Volcano plots were created to display the statistical significance (P-values) and effect sizes (beta value changes per age and the standardized coefficients for the age term) of each CpG site. Two types of Manhattan plots were used to show the genome-wide P-values and the standardized coefficients for each CpG site, organized by their chromosomal positions. Second, the age-related changes for each CpG site that exhibited a significant quadratic change pattern were visualized as heatmaps. Each raw beta value of the CpG sites was converted into a Z-value and categorized into two patterns: U-shaped and inverse U-shaped associations with age. These were sorted by the calculated vertex age of the quadratic curve and visualized as heatmaps, with higher Z-scores shown in red and lower Z-scores in blue. Additionally, examples of genes containing CpG sites with significant U-shaped or inverse U-shaped methylation patterns associated with age were provided. For this purpose, normalized heatmaps were created to further emphasize age-related changes, where the highest methylation level across age groups was set to 1, and the lowest methylation level was set to 0 for each CpG site (Figure 10).

### Results Demographics

Of the total of 10,420 samples, the age data were missing in 123 samples. The remaining 10,297 samples (98.8%) were used for the analysis. The dataset comprised of 5,156 males (50.1%) and 5,141 females (49.9%). The mean age of the sample was 53.9 years (SD = 14.1 years). Age distribution is shown in Figure 3.

**Figure 3.**
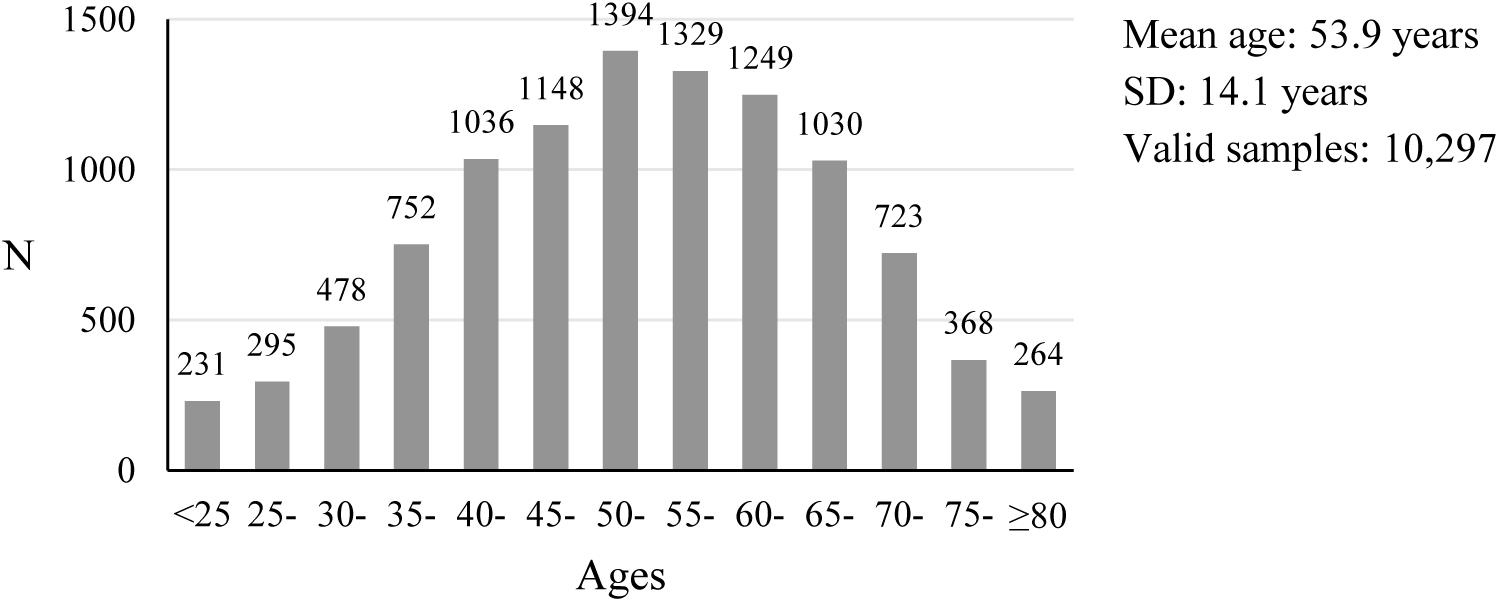
**Age composition**

### Linear regression analysis: the effect of age on DNA methylation

Among the 864,627 CpG sites, 278,704 CpG sites (32.2%) showed a significant linear association with age (P-value for age term of the regression < 5.7815×10^-8^). Significant increases in DNA methylation with age were observed in 72,480 CpG sites (8.4% of the total CpG sites, 26.0% of the significant age-associated CpG sites) and significant decreases of DNA methylation with age were observed in 206,224 CpG sites (23.9% of the total CpG sites, 74.0% of significant age-associated CpG sites). The histogram for the distribution of the regression coefficient and the standardized regression coefficient of the age term were shown in Figure 4.

**Figure 4.**
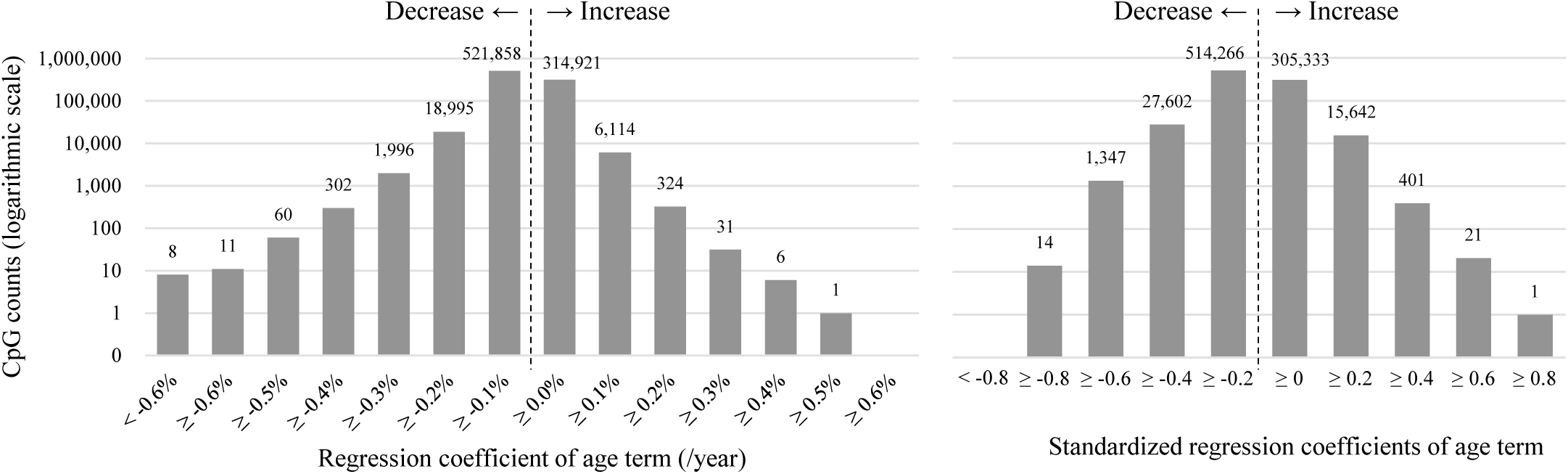
**Distribution of the coefficients**

The visualized plots (volcano plots and Manhattan plots) of the distribution of the relationships between the regression coefficients, the standardized regression coefficients, the location in chromosomes, and statistical significance (-log_10_P) were shown in Figure 5. ELOLV2: cg16867657, a well-known gene whose DNA methylation showed a significant association with age [20], was also detected in this dataset.

**Figure 5.**
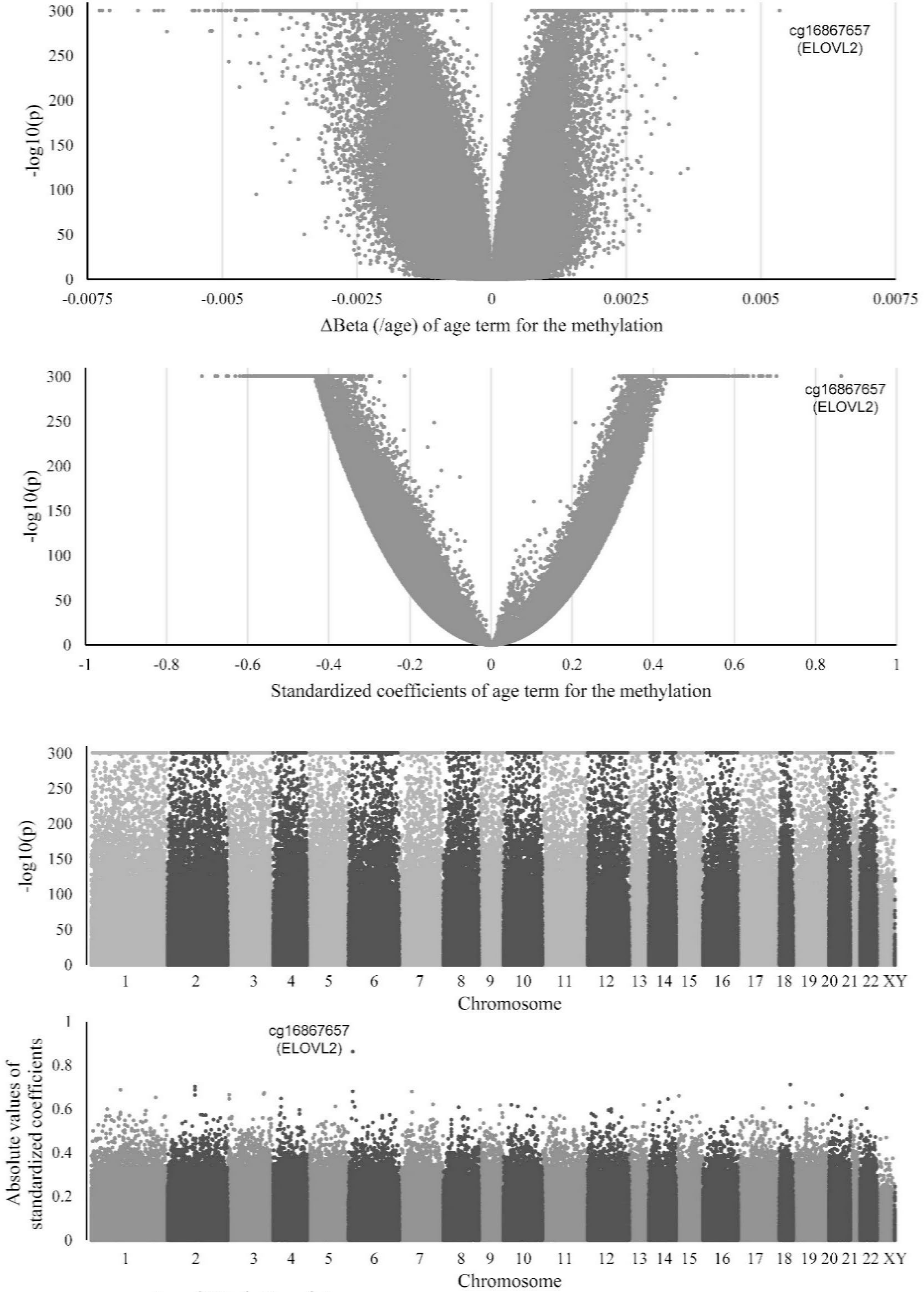
**Volcano plots and Manhattan Plots**

### Hierarchical regression analysis

Hierarchical multiple regression analysis incorporating an age-quadratic model identified 47,773 CpG sites with significantly increased explanatory power (P-value for ΔR² < 5.7815 × 10 ), indicating that the quadratic model provided a better fit for these associations. Consequently, as shown in Figure 6, 243,438 CpG sites exhibited only a significant linear association with age (P-value for ΔR² ≥ 5.7815 × 10 ) (A), 12,507 CpG sites showed significance only in the U-shaped or inverse U-shaped relationship with age (B), and 35,266 CpG sites exhibited both a significant linear association with age and were further explained by the quadratic model (C).

**Figure 6.**
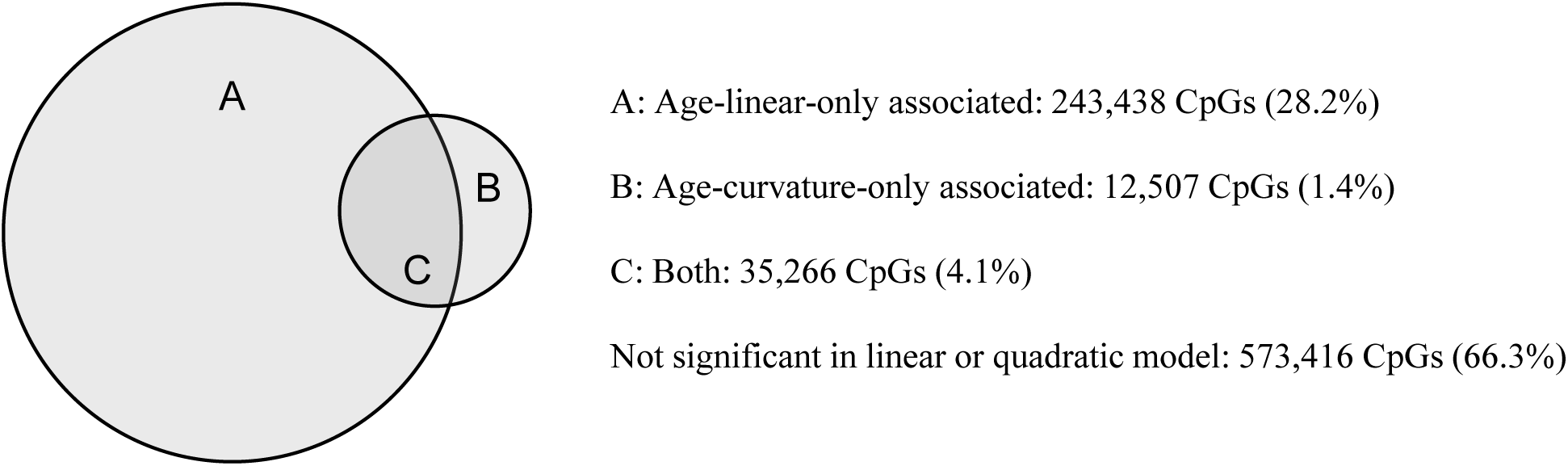
Distribution of Age-Associated CpGs Across Linear and Quadratic. Of the total 864,627 CpGs, 243,438 CpGs (28.2%) showed a significant linear association with age only, 12,506 CpGs (1.4%) showed a significant U-shaped or inverse U-shaped curve association with age only (B), and 35,265 CpGs (4.1%) exhibited both a significant linear association with age and were further explained by the quadratic model (C). The remaining 573,416 CpGs (66.3%) showed neither a linear nor a U-shaped or inverse U-shaped curve association with age.

As illustrated in Figure 7A, the standardized coefficients for the age term are more prevalent in the negative range, with the majority of CpG sites clustering between -0.2 and +0.2. Figure 7B shows the P-value distribution for the significance of the age term in the first-step regression analysis. Figure 7C illustrates the distribution of changes in R² values upon the addition of the age² term to the regression model and Figure 7D presents the density graph of P-values for the change in R².

**Figure 7.**
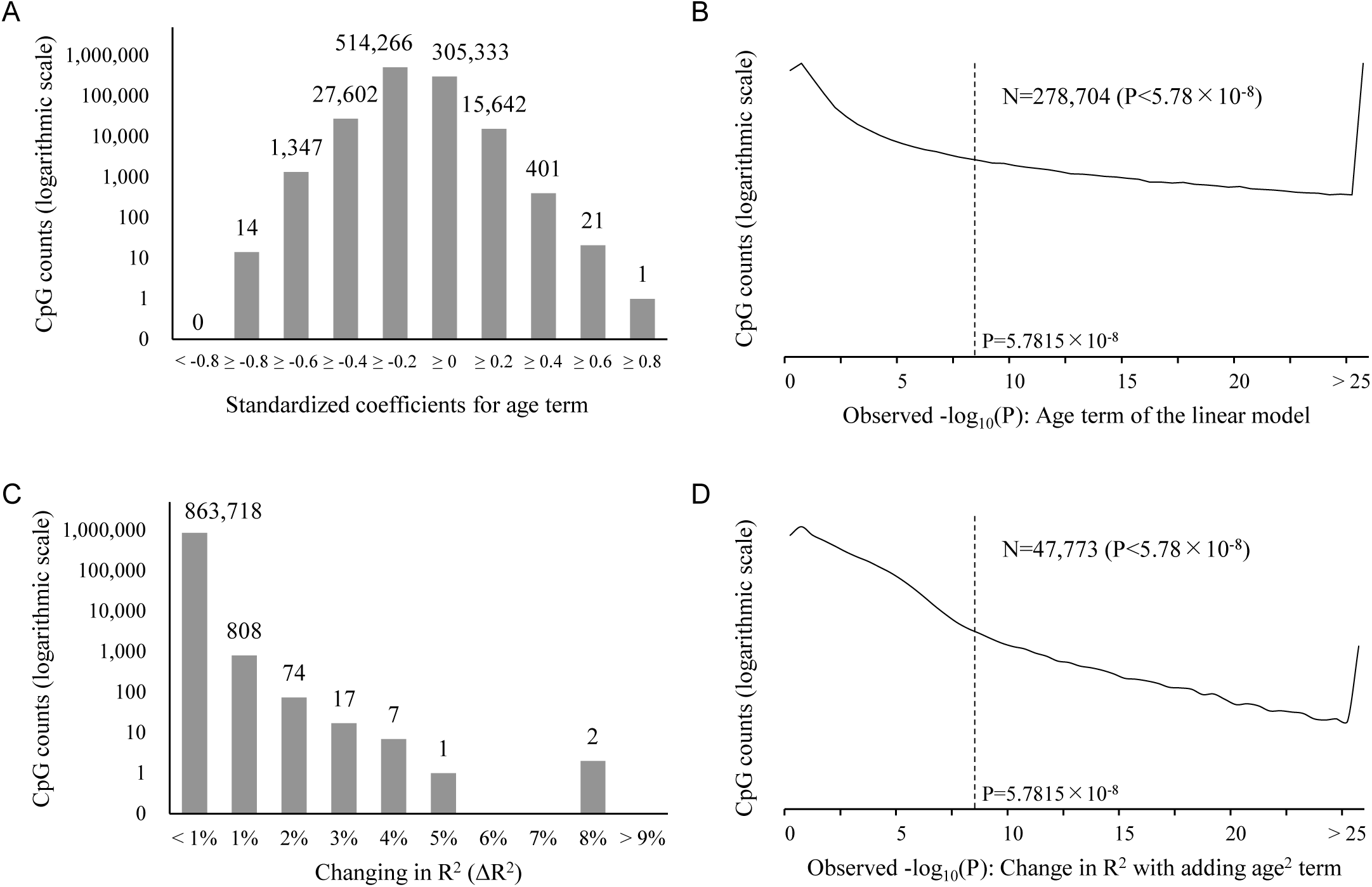
**Distribution of the regression results values** A: Histogram of the standardized coefficients for the age term. B: Density graph of P-values indicating the significance of the age term. C: Histogram showing the change in R² from adding the age² term to the regression. D: Density graph of P-values for the change in R² in the hierarchical multiple regression analyses.

### CpG sites with U-Shaped or Inverse U-Shaped Associations with Age

Of the 47,773 CpG sites identified as having a quadratic distribution of DNA methylation with age in the hierarchical multiple regression analysis, the trough or peak ages were calculated based on the regression results (Figure 2B). The distribution of these estimated trough or peak ages is shown in Figure 8. The CpG sites exhibiting U-shaped curves with age were less common in middle age and more prevalent in youth (< 30s) or old age (≥ 70s). In other words, for those with a U-shape association with age CpG sites, their troughs occur during youth or old age. This indicates that among CpG sites forming U-shaped DNA methylation patterns in relation to age, there are a small number of sites where DNA methylation levels decrease rapidly during early childhood and subsequently rise again with advancing age. Additionally, there are a larger number of sites where methylation levels consistently decline throughout the first half of life and begin to rise again in old age. On the other hand, the CpG sites exhibiting inverse U-shaped curves in the relationship between DNA methylation and age tended to have their methylation peaks more commonly during middle age. This means that, in humans, there are many CpG sites where DNA methylation levels are lower during youth but increase during middle age, then go down again during old age.

**Figure 8.**
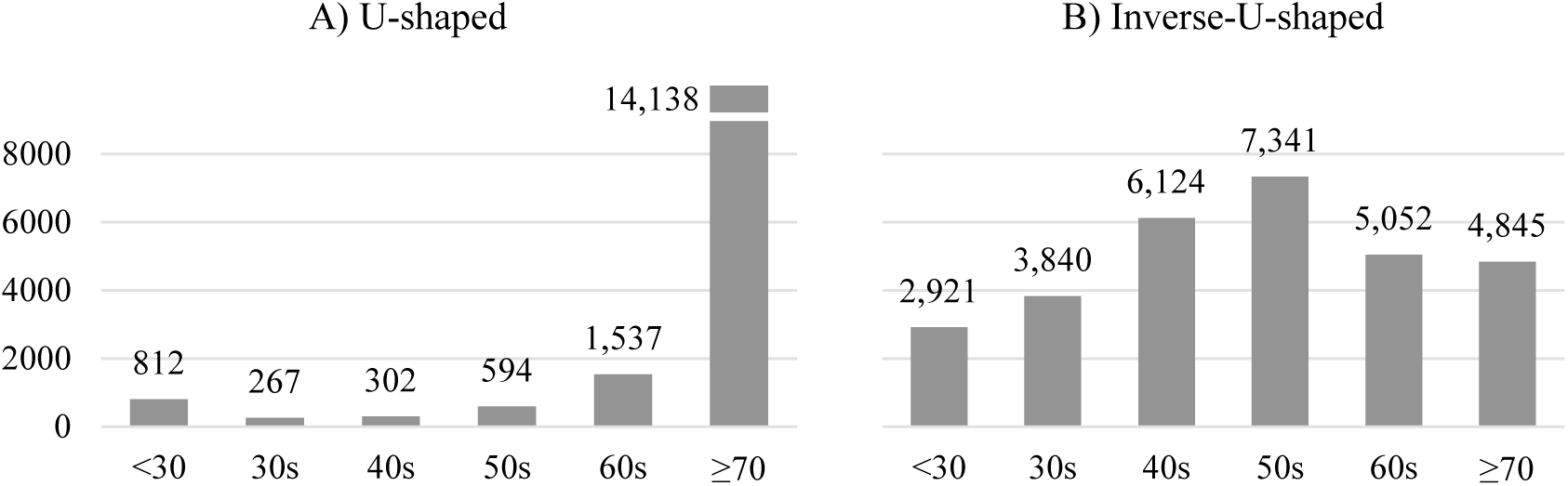
Distribution of estimated trough or peak ages. Estimated trough or peak ages of DNA methylation levels determined by the quadratic regression model. This distribution represents the peak or trough ages for 47,773 CpG sites where the change in ΔR² was significant in the hierarchical regression analysis.

The distribution of the age-specific mean DNA methylation values for each CpG site is shown in Figure 9. This figure illustrates how U-shaped or inverse U-shaped methylation changes occur with age. We visualized the trends in raw β-values, unadjusted for sex or WBC composition. Of the CpG sites showing significant U-shaped changes in DNA methylation with age, 2,700 CpG sites with vertex ages between 30 and 70 years are shown in Figure 9A, visualizing CpG sites that tend to show increased DNA methylation in young and old age, while DNA methylation decreases in middle age. Additionally, 22,357 CpG sites with significant inverse U-shaped changes and vertex ages between 30 and 70 years are shown in Figure 9B, categorized by decade into the 30s, 40s, 50s, and 60s, visualizing CpG sites that tend to show increased DNA methylation in middle age, while DNA methylation decreases in young and old age.

**Figure 9.**
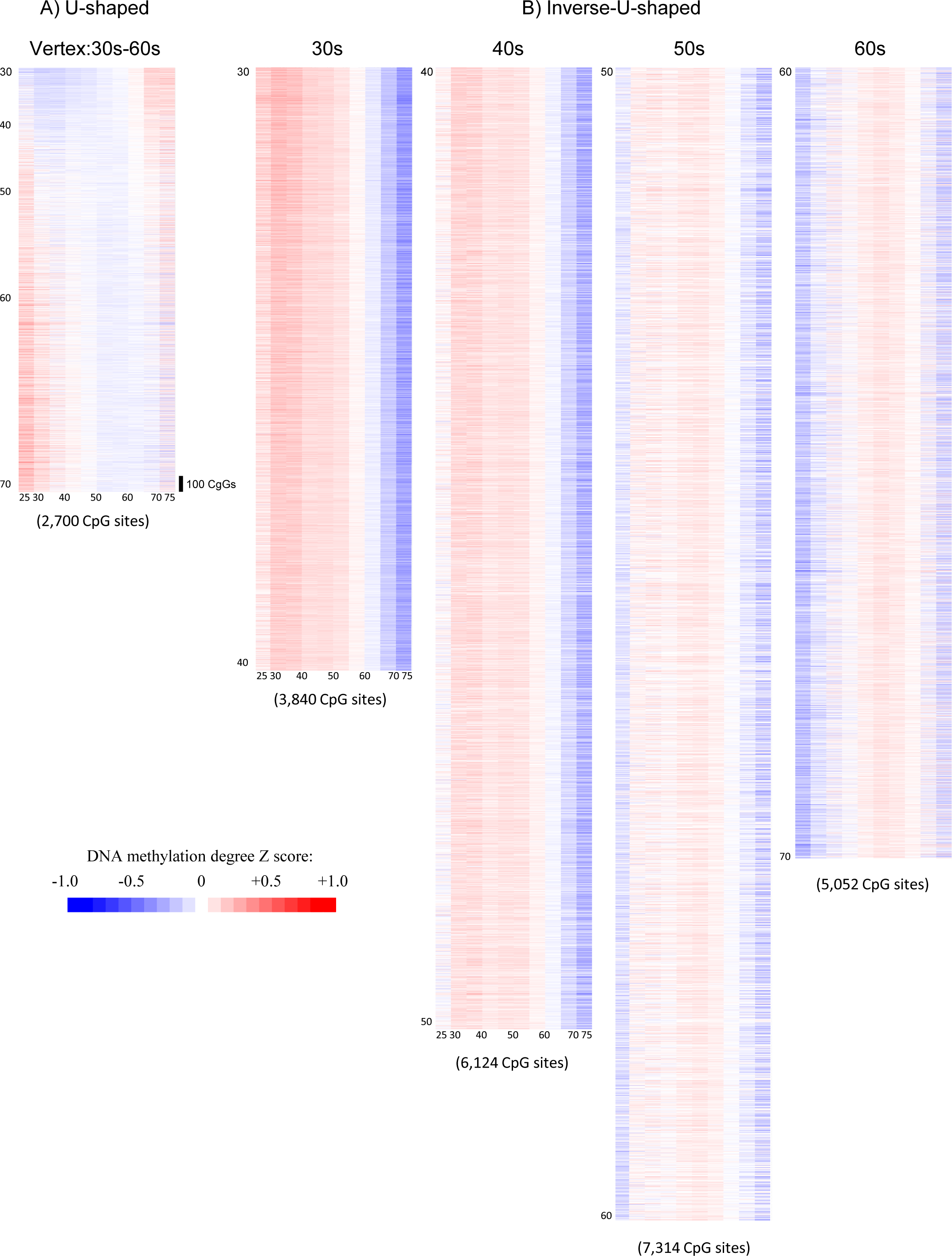
Heatmaps of quadratic pattern CpG sites. The temporal changes of 47,773 CpG sites, which exhibited a significant quadratic pattern in the hierarchical regression analysis, are visualized as a heatmap. Each color represents the change in Z-score (standardized value). The vertical axis represents each CpG site, while the horizontal axis shows the average DNAm raw β-values for each 5-year age group, unadjusted for sex or WBC composition. A) CpG sites with U-shaped associations with age are shown. B) CpG sites with inverse-U-shaped associations with age are shown.

**Figure 10.**
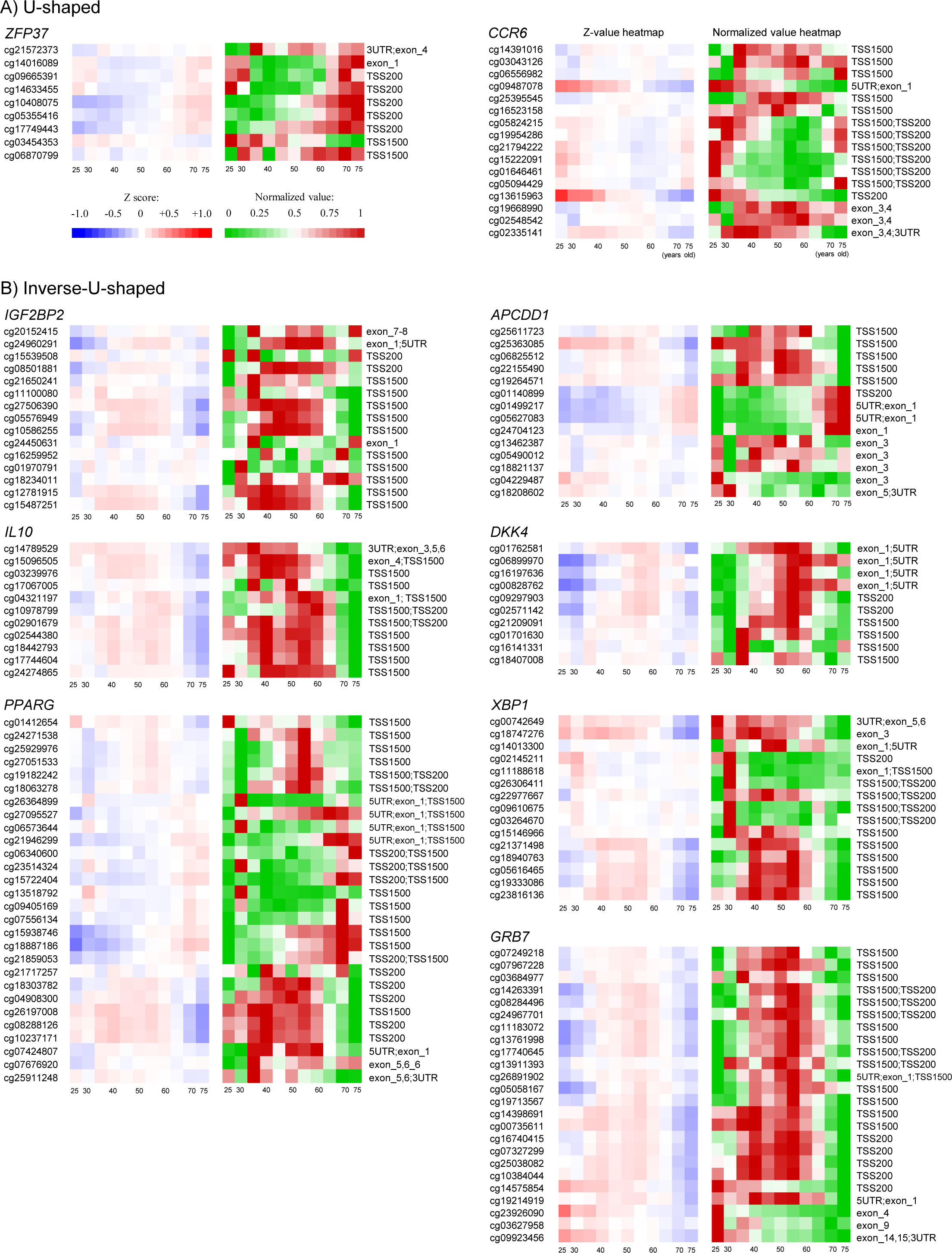
Example of the genes with CpG sites exhibiting quadratic patterns with age. Genes containing CpG sites with significant U-shaped or inverse U-shaped methylation patterns associated with age are exemplified. The left panel shows a heatmap of Z-values for each CpG site, while the right panel displays a heatmap of standardized values. CpG sites within each gene are sorted based on their chromosomal positions.

### Representative genes showing U-shaped or inverse U-shaped DNA methylation associations with age

To investigate which CpG sites of genes exhibit U-shaped or inverse U-shaped DNA methylation patterns associated with age, the frequency of CpG sites associated with specific genes was counted. Supplemental Table 1 lists CpG sites with a calculated vertex age value between 30 and 70 years that are annotated with known genes.

For CpG sites showing U-shaped curve associations, 16 genes were associated with CpG sites appearing at least twice. Among these, 10 genes had over 20% of their CpG sites showing significant U-shaped associations with age: *UQCRB, FUT7, SPATA13-AS1, LINC00426, IGFL2, ZBED2, CCR6, ZFP37, CPLANE2,* and *CD2*. Notable examples of U-

shaped DNA methylation patterns are shown in Figure 10A, highlighting *ZFP37*and *CCR6*. For *ZFP37*, significant U-shaped associations were found in the exon1 and TSS200 regions, with significant DNA methylation reductions around ages 40–50. For *CCR6*, significant U-shaped associations were observed in the TSS1500 and TSS200 regions, with a remarkable decrease in DNA methylation levels between the 40s and 60s concentrated in specific CpG sites.

For CpG sites showing inverse U-shaped curve associations, 703 genes were associated with CpG sites appearing at least twice. Among these, 189 genes had over 20% of their CpG sites showing significant inverse U-shaped associations with age. Examples of remarkable inverse U-shaped DNA methylation patterns are shown in Figure 10B, highlighting *IGF2BP2, APCDD1, IL10, DKK4, PPARG, XBP1,* and *GRB7*. In *IGF2BP2*, DNA methylation levels peak in the 40s–50s were observed in the TSS1500 region. In *APCDD1*, DNA methylation levels also peaked in the TSS1500 region during the 40s–50s. In *IL10*, DNA methylation levels peaked in the TSS1500 region from the late 30s to the early 60s. In *DKK4*, DNA methylation levels peaked around age 55 in the exon1 and TSS200 regions. In *PPARG*, peaks were observed in the TSS200 and TSS1500 regions between the late 30s and 50s, with another peak around age 55 in the same regions. For *XBP1*, DNA methylation levels peaked in the TSS1500 region during the 40s–50s. In *GRB7*, DNA methylation peaked across almost the entire gene, TSS200 and TSS1500 regions, with peaks showing from the 40s to the early 60s.

## Discussion

In this study, we analyzed a large-scale epigenome dataset from humans and demonstrated that changes in DNA methylation with age are not exclusively monotonic increases or decreases. Instead, we identified numerous CpG sites / genes that exhibit U-shaped or inverse U-shaped DNA methylation levels exhibiting troughs or peaks at specific ages. This finding suggests that humans do not merely experience monotonic changes in gene expression with age. Rather, multiple genes are either upregulated or downregulated at specific stages of life, likely to enable certain physiological processes tailored to each life stage. Furthermore, this result strongly suggests that age-related changes in DNA methylation are not merely induced by passive, physicochemical wear and tear over time [5, 6] but are likely subject to some form of active regulation.

A substantial number of genes were identified to exhibit quadratic associations with age. While a comprehensive discussion of all these genes is beyond the scope of this study, some of the previously highlighted genes suggest potential pathological or physiological significance. DNA methylation in the TSS200-TS1500 promoter region is known to have a significant impact on gene expression [21, 22]. Notably, in the genes highlighted in this study, significant age-related U-shaped or inverse U-shaped patterns were detected in CpG sites within that region. This finding suggests the existence of gene regulatory mechanisms that are tightly connected with specific life stages. For instance, *CCR6*, which exhibited a DNA methylation trough between the ages of 40 and 60, particularly around the 50s, is well-recognized as a gene implicated in the pathogenesis of rheumatoid arthritis [23, 24]. From the perspective of DNA methylation, this may indicate that the epigenetic status leading to *CCR6* expression peaks between the ages of 40 and 60.

Interestingly, while the prevalence of rheumatoid arthritis increases with age, the peak age of onset aligns with this range [25]. Another example is the *PPAR*γ gene. The downregulation of *PPAR*γ is known to contribute to metabolic abnormalities and the development of type 2 diabetes [26] . In this study, DNA methylation at *PPAR*γ was found to be significantly elevated between the ages of 35 and 60, suggesting that its expression may be most suppressed during middle age. This could potentially relate to the increased prevalence of obesity and metabolic syndrome observed during this period. Furthermore, *GBR7* exhibited a unique pattern of DNA methylation across its TSS200 to TSS1500 regions, with strong DNA methylation starting around the age of 40, a pronounced peak in the 50s, and subsequent DNA methylation decrease beginning in the 60s. While *GBR7* expression is associated with breast cancer progression [27], it has also been implicated to contribute to normal lactation [28]. It is possible that *GBR7* expression is maintained during the reproductive years to support lactation, while its suppression after the programmed end of the lactation period around the age of 40 may serve to prevent breast cancer. However, the physiological mechanism of the observed “re-demethylation” during old age in this gene remain unclear.

Our results align partially with previous studies that identified age-associated differentially methylated regions and characterized the phenomenon of epigenetic drift. For instance, prior research reported genome-wide hypomethylation alongside promoter hypermethylation with aging [4,5]. A majority of the results of this study aligns with these observations. However, while other studies primarily assumed linear trends, our data reveal that a significant subset of CpG sites follows non-linear trajectories, indicating that age-related DNA methylation changes are more dynamic and multifaceted than previously understood. This novel observation calls for a re-evaluation of the conventional frameworks used in epigenome-wide association studies (EWAS) and epigenetic age studies, particularly those relying solely on linear modeling. Existing epigenetic age estimation methods or epigenome clocks are calculated based on methylation status—simply whether specific CpG sites are hypermethylated or hypomethylated. Many studies use only linear methylation changes to estimate epigenetic age and consider it interchangeable with biological age. In these models, an epigenome clock age higher than chronological age is typically interpreted as representing accelerated biological decline, implying that such changes should be avoided. These investigations also sometimes claim that reversing the epigenome clock age equates to rejuvenation and is inherently beneficial. Our study suggests that the regulation of the human body is complex, with distinct gene expression switches turning on or off at specific life stages. This highlights the need to critically evaluate whether the genes investigated in the studies of epigenomics truly expressed linearly with aging. For example, it is important to consider that if anti-aging interventions were to cause an increase or decrease in methylation at specific genes, the interpretation of such changes could depend on the potential non-linear relationship between age and their DNA methylation levels. For genes with non-linear associations, the meaning of these changes could reverse depending on the individual’s age, raising the question of whether such changes represent aging acceleration or anti-aging effects.

Future studies should explore these issues and focus on determining the sequence of DNA methylation peaks or troughs among the genes identified in this study. Investigating whether these changes occur in a cascade-like manner will be particularly valuable. Moreover, it is worth examining whether “middle-age markers” based on elevated or reduced DNA methylation levels in specific gene groups or epigenome age estimation methods using quadratic models can be developed. Such analyses could open new avenues in understanding the relationship between epigenomics and aging.

This study has certain limitations. The non-clinical population samples used in this analysis were not collected through random sampling of the general population, which may introduce biases related to socio-economic factors or literacy. Additionally, racial and ethnic differences were not considered or adjusted for, and therefore, the generalizability of these findings to the broader population cannot be assured.

## Conclusion

Among the 864,627 CpG sites analyzed, 8.4% showed a significant increase in DNA methylation with age, 23.9% showed a significant decrease, and 5.5% were significantly better explained by a quadratic model. Within this subset, many CpG sites exhibited an inverse U-shaped relationship with age, with methylation levels peaking during middle age. Numerous genes showing significant U-shaped or inverse U-shaped associations were identified in relation to diseases known to have peak onset ages during middle adulthood. These findings suggest that the relationship between DNA methylation and aging is not simply linear but more dynamic, with gene expression being promoted or suppressed at specific ages or life stages related to age-dependent processes.

## CRediT authorship contribution statement

AS: Conceptualization, Formal analysis, Investigation, Methodology, Resources, Software, Writing – original draft. KY: Validation, Writing – review & editing. TI: Validation, Writing – review & editing. TS: Validation, Writing – review & editing. TN: Validation, Writing – review & editing. BA: Validation, Writing – review & editing. TS: Writing – original draft, Writing – review & editing. VD: Data curation, Resources, Writing – review & editing. RS: Data curation, Supervision, Writing – review & editing. GS: Conceptualization, Data curation, Funding acquisition, Investigation, Project administration, Resources, Supervision, Writing – review & editing.

## Data availability

The data that support the findings of this study are not publicly available due to protection of patient data in accordance with maintaining HIPAA compliance. However, the data can be made available upon reasonable request after signing a Data Use Agreement.

## Acknowledgments

The dataset was provided free of charge by TruDiagnostic Inc. for this investigation. No external funding was received for this study.

## Gene Symbols and Full Names

Symbol Full: gene name
ELOVL2: ELongation of very long chain fatty acids protein 2
ZFP37: Zinc Finger Protein 37
CCR6 C-C: Motif Chemokine Receptor 6
IGF2BP2: Insulin Like Growth Factor 2 MRNA Binding Protein 2
APCDD1: APC Down-Regulated 1
IL10: Interleukin 10
DKK4: Dickkopf WNT Signaling Pathway Inhibitor 4
PPARG: Peroxisome Proliferator Activated Receptor Gamma
XBP1: X-Box Binding Protein 1
GRB7: Growth Factor Receptor Bound Protein 7

